# Characterizing sensitivity and coverage of clinical WGS as a diagnostic test for genetic disorders

**DOI:** 10.1101/2020.04.01.019570

**Authors:** Yan Sun, Fengxia Liu, Chunna Fan, Yaoshen Wang, Lijie Song, Zhonghai Fang, Rui Han, Zhonghua Wang, Xiaodan Wang, Ziying Yang, Zhenpeng Xu, Jiguang Peng, Chaonan Shi, Hongyun Zhang, Wei Dong, Hui Huang, Yun Li, Yanqun Le, Jun Sun, Zhiyu Peng

**Author notes:** Correspondence: Jun Sun; Zhiyu Peng. Yan Sun, Fengxia Liu, Chunna Fan and Yaoshen Wang contributed equally to this work.

## Abstract

**Background:** With the reduce of cost and incomparable advantages, WGS is likely to change the way of clinical diagnosis of rare and undiagnosed diseases. However, the sensitivity and breadth of coverage of clinical WGS as a diagnostic test for genetic disorders has not been fully evaluated, especially for CNV detection.

**Methods:** All the samples underwent WGS using MGISEQ-2000. The performance of NA12878, YH cell line, and the Chinese trios were measured for sensitivity, PPV, depth and breadth of coverage. We also compared the performance of WES and WGS using NA12878. The sensitivity and PPV were tested using family-based trio design in the Chinese trios. We also developed a systematic WGS pipeline for the analysis of 8 clinical cases with known disease-causing variants.

**Results:** In general, the sensitivity and PPV for SNV/indel detection increased with mean depth, and reached a plateau at a ~40X mean depth using down-sampling samples of NA12878. With a mean depth of 40X, the sensitivity of homozyous and heterozygous SNPs of NA12878 was >99.25% and >99.50% respectively, and PPVs were 99.97% and 98.96%. Homozygous and heterozygous indels showed lower sensitivity and PPV. The sensitivity and PPV is still not 100% even with a mean depth of ~150X. We also observed a substantial variation in the sensitivity of CNV detection across different tools, especially in CNVs with a size of less than 1kb. In general, the breadth of coverage for disease-associated genes and CNVs increased with mean depth. The sensitivity and coverage of WGS (~40X) is better than WES (~120X). Among the Chinese trios with ~40X mean depth, the sensitivity in the offspring was >99.48% and >96.36% for SNP and indel detection, and PPVs were 99.86% and 97.93%. All the 12 previously validated variants in the 8 clinical cases were successfully detected by our WGS pipeline.

**Conclusions:** The current standard of a mean depth of 40X may be sufficient for SNV/indel detection and identification of most CNVs. Clinical scientists should know the range of sensitivity and PPV for different classes of variants for a particular WGS pipeline, which would be useful when interpreting and delivering clinical reports.

## Background

So far, there are more than 8,000 Mendelian diseases recorded by OMIM (Online Mendelian Inheritance in Man), more than 5,000 of which have phenotype description and molecular basis. The rapid development of massively parallel sequencing (MPS) technology has revolutionized the field of genetic diagnosis in clinical setting, making it possible for MPS to be a routine part of clinical care. Recently, whole-exome sequencing (WES) and whole genome sequencing (WGS) are routinely used and are gradually being optimized for the detection of rare and common genetic variants in humans [1–4]. Comparing to WES, WGS is more powerful for detecting variants. Theatrically, WGS has the potential to identify nearly all forms of genetic variation [5], including single-nucleotide variants (SNVs) in both the protein-coding and noncoding regions (such as introns and promoters) of the genome, small insertions/deletions (indels), and copy-number variants (CNVs) [6, 7]. Without target region selection, WGS could provide a more uniform DP for the genome, making the detection of CNVs readier. What is more, WGS could provide higher sensitivity and higher yield in variant detection in the coding regions [8–10]. Several studies have demonstrated the advantages of WGS for variant detection [11–14]. For patients who highly suspected with genetic disorder, WGS maybe one and the optimal for further evaluation when the patient remains undiagnosed after clinical WES. With the reduce of the cost in sequencing and its incomparable advantages, WGS is likely to change the way of clinical diagnosis of rare and undiagnosed diseases and is bound to become a routine part of clinical care in the near future.

So far, WGS still follows the three steps since the invention of MPS technology: preparation of template, library construction and sequencing, and data analysis. Factors that influence the three steps (such as quality of genomic DNA [15, 16], library preparation [17–19], sequencing platform [18, 20], bioinformatics analytical pipeline [21]) may influence the results of clinical WGS. Building a standardized wet laboratory and bioinformatics pipeline can improve comparability of WGS. The analysis of a clinical WGS usually starts from quality evaluation. Mean DP is often recognized as a general indicator of overall sensitivity. It has been reported that variant calling is more reliable with increasing DP [22]. As a crucial factor in data quality, most centers conducting MPS technology would determine thresholds for average DP across WES/WGS, and determine the minimum DP that must be achieved for a certain fraction of target bases [23]. The breadth of coverage describes the fraction of the total target genomic region that is sequenced to an adequate depth in a particular assay [4]. The American College of Medical Genetics and Genomics (ACMG) recommends that 90%-95% breadth of coverage above a minimum threshold of 10X should be achieved for exome data with an average depth of 100X [23]. So far, a mean depth of 30-50X is the most widely used mean DP for WGS [24, 25]. However, the sensitivity and breadth of coverage of clinical WGS has not been fully evaluated, especially for CNV detection, some of which are associated with human disease [26–29]. For clinical WGS, the sensitivity and coverage of CNVs has not been comprehensively investigated.

In this study, we performed a systematic analysis of the sensitivity and coverage of clinical WGS using 5 gold standard samples (NA12878, YH, NA24631, NA24694 and NA24695). Then we applied clinical WGS for the reanalysis of 8 clinical cases with known disease-causing variants. The results may provide a reference for the laboratories who perform clinical WGS.

## Methods

### Study design and sequencing of samples

In order to test the sensitivity and coverage of clinical WGS, Genome in a Bottle (GIAB) sample HG001 (NA12878), HG005 (NA24631)/HG006 (NA24694)/HG007 (NA24695) (known as the Chinese son/father/mother), and YH cell line (a human lymphoblastoid cell line from first Asian genome donor) [30] were collected and sequenced using MGISEQ-2000 platform. All the samples used in this study are listed in Table 1.

Figure 1 shows the overall design of this study. First, in order to assess the recommended depth for proband-only WGS in clinical diagnostics, the analysis of the sensitivity and positive predictive value (PPV) of high-confidence SNPs/indels, the sensitivity of CNV detection, and the depth and breadth of coverage for disease-associated genes and CNVs were performed using down-sampling samples of NA12878-1. The results of sensitivity and PPV from a single genome may be difficult to generalize to a range of samples [31]. So in this part, similar analyses were also performed for down-sampling samples of another high depth sequencing sample of YH. After determining the recommended mean DP for proband-only WGS, the Chinese trios (NA24631, NA24694 and NA24695) with recommended mean DP were used to test the sensitivity and PPV of trio-based WGS when taking advantage of the family-based trio design in clinical WGS. Using down-sampling samples of NA12878-1 and NA12878-2, we also compared the performance of WES and WGS using the recommended mean DP. Finally, we analyzed 8 clinical cases with known disease causing variants using our WGS pipeline (Figure 1).

**Figure 1.**
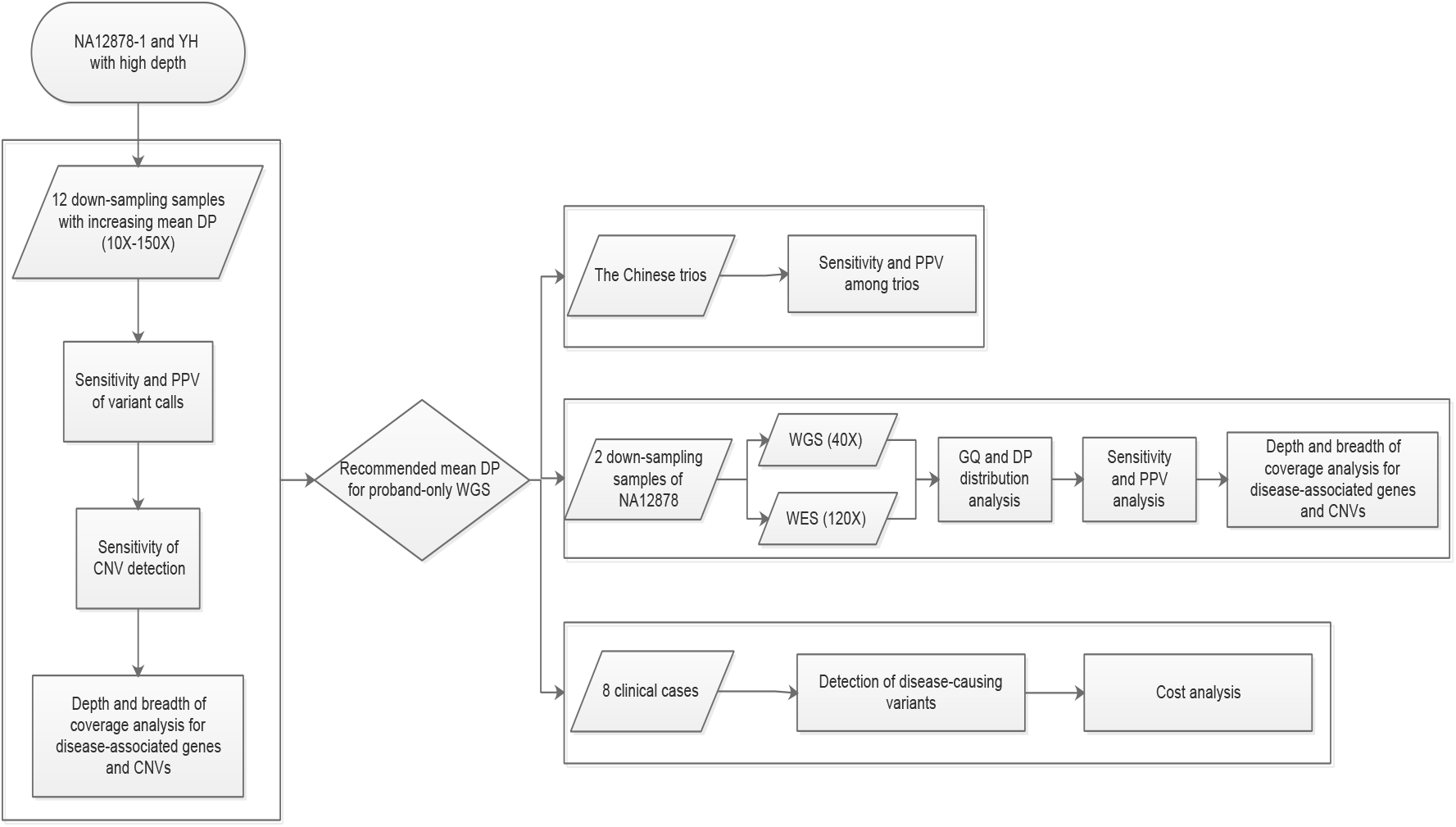
Study design

Based on the study design, DNA samples of NA12878, NA24631, NA24694 and NA24695 were procured from Coriell (Camden, NJ). 1 μg of DNA were used for YH and the GIAB Chinese family to generate paired-end reads of 100 bp. The most widely used NA12878 were sequenced 2 times (WES and WGS) (Table 1). This study and all the protocols were approved by the ethics committee of BGI (NO. BGI-IRB19143).

### Alignment and variant calling

In this study, a standard bioinformatics pipeline was used for the analysis of all the samples (Supplementary material). In order to eliminate variability due to differences by various bioinformatics tools, we developed a standard bioinformatics pipeline, which included current widely used tools (Supplementary material). In general, after removal of sequencing adapters and low-quality reads, “clean reads” were aligned to the GRCh37 with BWA 0.7.12-r1039 [32]. Genome Analysis Toolkit (GATK)-package-4.0.11.0 [33] MarkDuplicates was used to remove duplicate reads. After realignment around indels and quality scores re-calibration using GATK-package-4.0.11.0 [33], VCF files were then generated using GATK for further analysis. As for the CNV detection, 3 widely used tools (CNVnator [34], BreakDancer [35] and LUMPY [36]) were performed in this study.

After trimming sequencing adapters and consecutive low-quality bases using fastp [37], the clean reads of NA12878-1 were down-sampled by the sequencing depth of 10X (NA12878-1_10X), 20X (NA12878-1_20X), 30X (NA12878-1_30X), 40X (NA12878-1_40X), 50X (NA12878-1_50X), 60X NA12878-1_60X), 70X (NA12878-1_70X), 80X (NA12878-1_80X), 90X (NA12878-1_90X), 100X (NA12878-1_100X), 120X (NA12878-1_120X) and 150X (NA12878-1_150X) using seqtk (https://github.com/lh3/seqtk). The high coverage NA12878-1 sample was sequenced using MGISEQ-2000 platform to an average depth of ~197X (PE100) using PCR-based library construction. The clean reads of YH were also down-sampled by sequencing depth.

### Sensitivity and positive predictive value of variant calls

To evaluate the performance of variant calls in detecting true genotypes, the high-confidence calls (SNPs and indels) for GIAB sample HG001 (NA12878) and HG005/HG006/HG007 (Chinese son/father/mother) were considered as true-positive calls (v3.3.2) [38]. We restricted the calculation of sensitivity (high-confidence calls detected by our method/all the high-confidence calls in GIAB) and PPV (high-confidence calls detected by our method/all the variants detected by our method) of variant calls to the high confidence region (v3.3.2) [38]. High-confidence variant calls and regions tend to include a subset of variants and regions that are easier to detect [38]. The sensitivity and PPV of variant calls in YH sample were also evaluated. The alleles validated by the Illumina 1M BeadChip were considered as “true-positive” calls for YH [30]. Genotype quality (GQ) and DP were used to filter out variants with erroneous variant calls.

### Breadth of coverage for disease-associated genes and CNVs

In order to assess the recommended depth for proband-only WGS in clinical diagnostics, we collected a total of 6 gene sets for coverage analysis of disease-associated genes: ACMG59 [39], ClinVar [40] (3,824 genes, accessed on 19 February 2019), Genetic Home Reference [41] (1,471 genes, accessed on 2 July 2019), HGMD [42] (8,171 genes, professional March 2018), OMIM [43] (3,835 genes, accessed on 4 April 2018) and Orphanet [44] (2,405 genes, accessed on 2 July 2019). For the annotation of gene regions, we used the information of NCBI annotation release 104. For different transcripts, we first used the transcripts used in the HGMD database. The rest of the gene region was a combination of the regions of all transcripts. Coverage analysis of the 12 down-sampling samples of NA12878-1 and YH for the 6 gene sets were performed for evaluation.

Genes in a single gene set of the 6 gene sets is incomplete. In order to generalize a new gene list that contains all the putative disease-associated genes, we compiled a list of 8,394 putative disease-associated genes from the 6 gene sets (Supplementary Table 1). This new gene set was generated using the following criteria: 1) a gene was kept when it was recorded with association status of “assessed” in the Orphanet database, and with one of the following association type: a) “Disease-causing germline mutation(s) in”, b) “Disease-causing germline mutation(s) (gain of function) in”, c) “Disease-causing germline mutation(s) (loss of function) in”, d) “Major susceptibility factor in”, e) “Modifying germline mutation in”, f) “Role in the phenotype of’, g) “Candidate gene tested in”; 2) a gene was kept when it was recorded as a disease related gene in the Genetic Home Reference database; 3) a gene was kept when it was recorded as a disease related gene and the molecular basis of the disease is known, unless the inheritance of the disease was recorded as “SMu” (somatic mutation) only in OMIM database; 4) a gene was kept when at least one variant of the gene was recorded as probable/possible pathological mutations in the HGMD database; 5) a gene was kept when at least one variant of the gene was recorded as pathogenic or likely pathogenic in the ClinVar database; 6) a gene was kept unless the gene was not recorded with well-defined gene ID or genome coordinate information in NCBI annotation release 104. This new gene list was used in the comparison of WES and WGS and is ideal for coverage analysis of clinical WGS.

We also performed coverage analysis of the 12 down-sampling samples of NA12878-1 and YH for CNVs described in the DECIPHER database (GRCh37_v9.29). Detailed information can be found in Supplementary Table 2.

### CNV analysis

Unlike SNP and indel, there are no perfect “gold standard” CNV dataset for benchmarking. In this study, in order to assess the recommended depth for proband-only WGS in clinical diagnostics, we conducted the evaluation of sensitivity of CNV detection using 3 CNV call sets of NA12878 from published papers [45–47]. CNV call set 1 was also used by Haraksingh et al [45] for the benchmarking of CNV detection from 17 commercially available arrays and low-coverage WGS. CNV call set 2 was called by a machine learning based approach (svclassify), and obtained from the GIAB Consortium benchmark SV calls resource [46]. CNV call set 3 included 874 deletions detected by both reference-based (a custom pipeline and PBHoney using both raw and error-corrected reads) and assembly-based analysis via single-molecule technologies [47]. All these 3 CNV call sets have been previously compiled from NA12878 for benchmarking and downloaded for further analysis in current study (Supplementary Table 3).

In the 12 down-sampling samples of NA12878-1, CNVnator (v0.3.2) [34], BreakDancer (v1.4.5) [35] and LUMPY (v0.2.13) [36] were assessed for sensitivity of CNV detection with default or recommended parameters. In a CNV call set, true positives were classified as CNVs with at least a 50% reciprocal overlap with CNVs in the call set (BEDTools) [48]. For benchmarking, we determined the number of gold standard CNVs detected in the 12 down-sampling samples of NA12878-1 for the 3 CNV call sets.

### Sensitivity and PPV of variant calls in the Chinese trios

To test the sensitivity and PPV of variant calls for trio-based WGS, we investigated the sensitivity and PPV when taking advantage of the family-based trio information in the Chinese trios. Using the segregation pattern, we focused on the autosomes and X chromosome of NA24631, NA24694 and NA24695. Taking advantage of the family-based trio design, we analyzed all variants (DP >= 10X) of both parents where one parent was consistently called as homozygous for the reference allele and the other as homozygous for the alt allele. We then used the variant calls (SNPs and indels) in the offspring to test the sensitivity and PPV in these loci.

## Results

### Sensitivity and positive predictive value of variant calls

Mean DP was recognized as a general indicator of overall sensitivity for SNV/indel detection [4]. In order to reveal the sensitivity of proband-only WGS for SNP/indel detection, 12 down-sampling samples of NA12878-1 with increasing of mean DP (10X-150X) were evaluated. For the 12 down-sampling samples of NA12878-1, we restricted the calculation of sensitivity and PPV of SNPs/indels to the high confidence region (v3.3.2). GQ (>= 20) and DP (>= 10) were used to filter out variants with erroneous variant calls. As a result, when the mean depth is more than 40X, the sensitivity of detecting homozygous and heterozygous SNPs is more than 99.25% and 99.50%, the sensitivity of homozygous and heterozygous indels is more than 88.50% and 89.09% respectively. The PPV (high confidence region) for homozygous and heterozygous SNPs exceeded 99.97% and 98.96%, the PPV for homozygous and heterozygous indels exceeded 98.93% and 84.26% respectively. Heterozygous indels showed the lowest PPV. Considering the sensitivity of SNP/indel detection and sequencing costs, a sequencing depth of ~40X provided the best value for SNP/indel detection, as is indicated by the trends in the sensitivity results (Figure 2).

**Figure 2.**
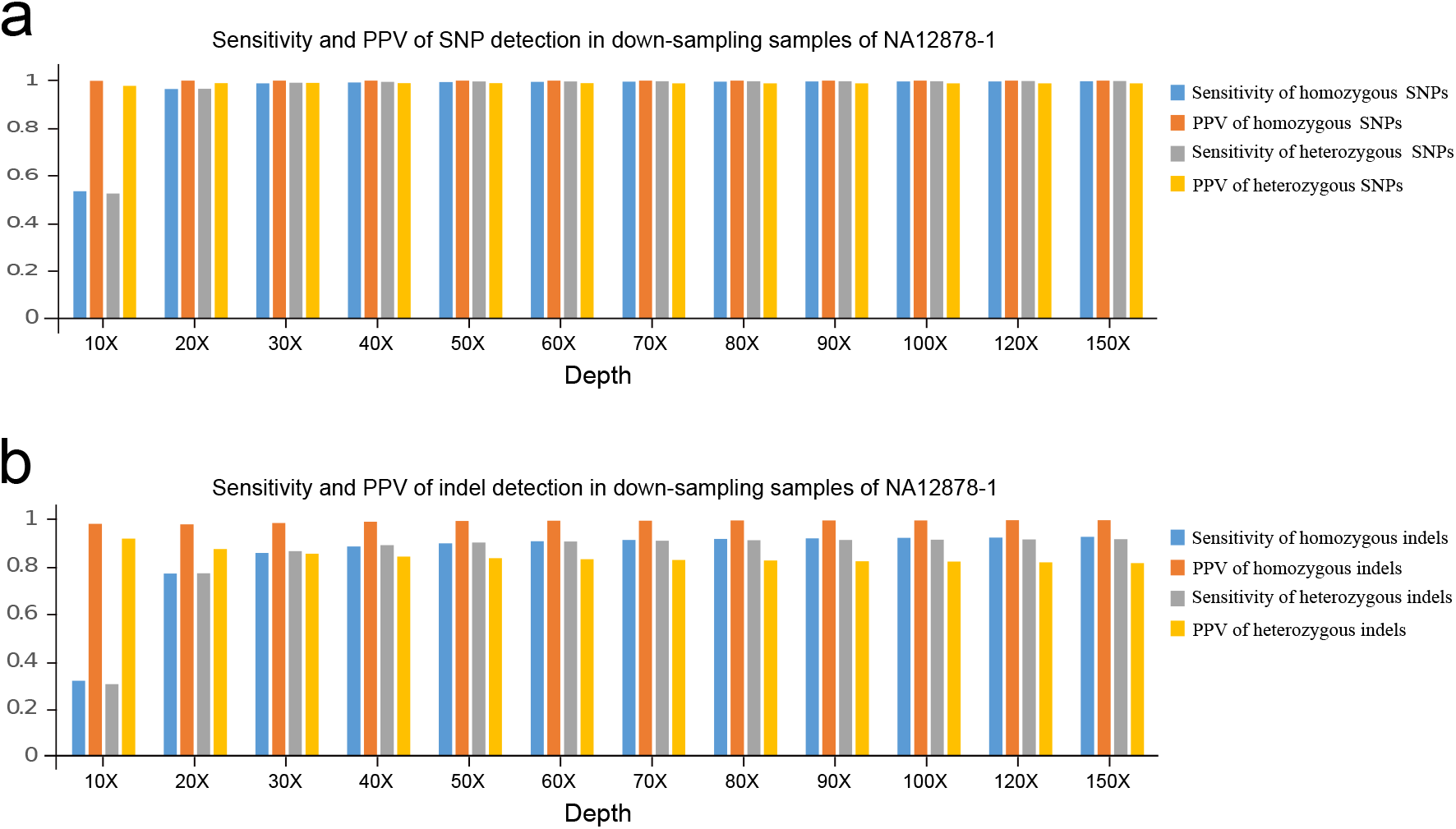
Sensitivity and PPV of variant calls from 12 down-sampling samples of NA12878-1

The sensitivity of homozygous and heterozygous SNPs exceeded 96.48% and 96.59%, and reached a plateau even with a mean depth of only 20X. The sensitivity significantly increased with sequencing depth from 10X to 30X for both homozygous and heterozygous indels, but reached a plateau at a ~40X mean depth. A mean depth of 40X could provide a percentage of more than 99.05% for sites covering more than 20X in the high confidence region. However, clinical scientist should know that, even with a DP of ~150X, the sensitivity and PPV is still not 100%. With a DP of 150X, the sensitivity for homozygous and heterozygous SNPs was 99.70% and 99.81%, the sensitivity of homozygous and heterozygous indels reached 92.57% and 91.57% respectively. With a DP of 150X, the PPV for homozygous and heterozygous SNPs was 99.97% and 98.82%, the PPV for homozygous and heterozygous indels reached 99.61% and 81.42% respectively.

The results of sensitivity and PPV from a single genome may be difficult to generalize to a range of samples [31]. So in this study, we also performed sensitivity and PPV analysis of SNPs (in the 1M validated region) for down-sampling samples of another high depth sequencing sample of YH (Supplementary Table 4). Similar results were obtained.

### Sensitivity of CNV detection

In order to detect the sensitivity of proband-only WGS for CNV detection. The 12 down-sampling samples of NA12878-1 (10X-150X) with increasing mean DP were evaluated. CNVnator (read depth) [34], BreakDancer (read pair) [35] and LUMPY (read depth and read pair) [36] were used for the detection of CNVs for the 12 down-sampling samples. In this study, a total of 3 CNV call datasets were assessed. The overall sensitivity of CNVnator, BreakDancer and LUMPY of the 12 down-sampling samples for CNV call set 1 (BreakDancer only included the detection of deletions in CNV call set 1), CNV call set 2 and CNV call set 3 was shown in Figure 3. In general, CNV calling is reliable with increasing DP (Figure 3a-c). At increasing sequencing depths, the trends of the sensitivity curves for the 3 CNV tools were different from one another. CNVnator showed a wide range in the sensitivity with varying DP, and the sensitivity visibly increased with mean depth, which indicates that the sensitivity of CNV detection is positively correlated with sequencing depth.

**Figure 3.**
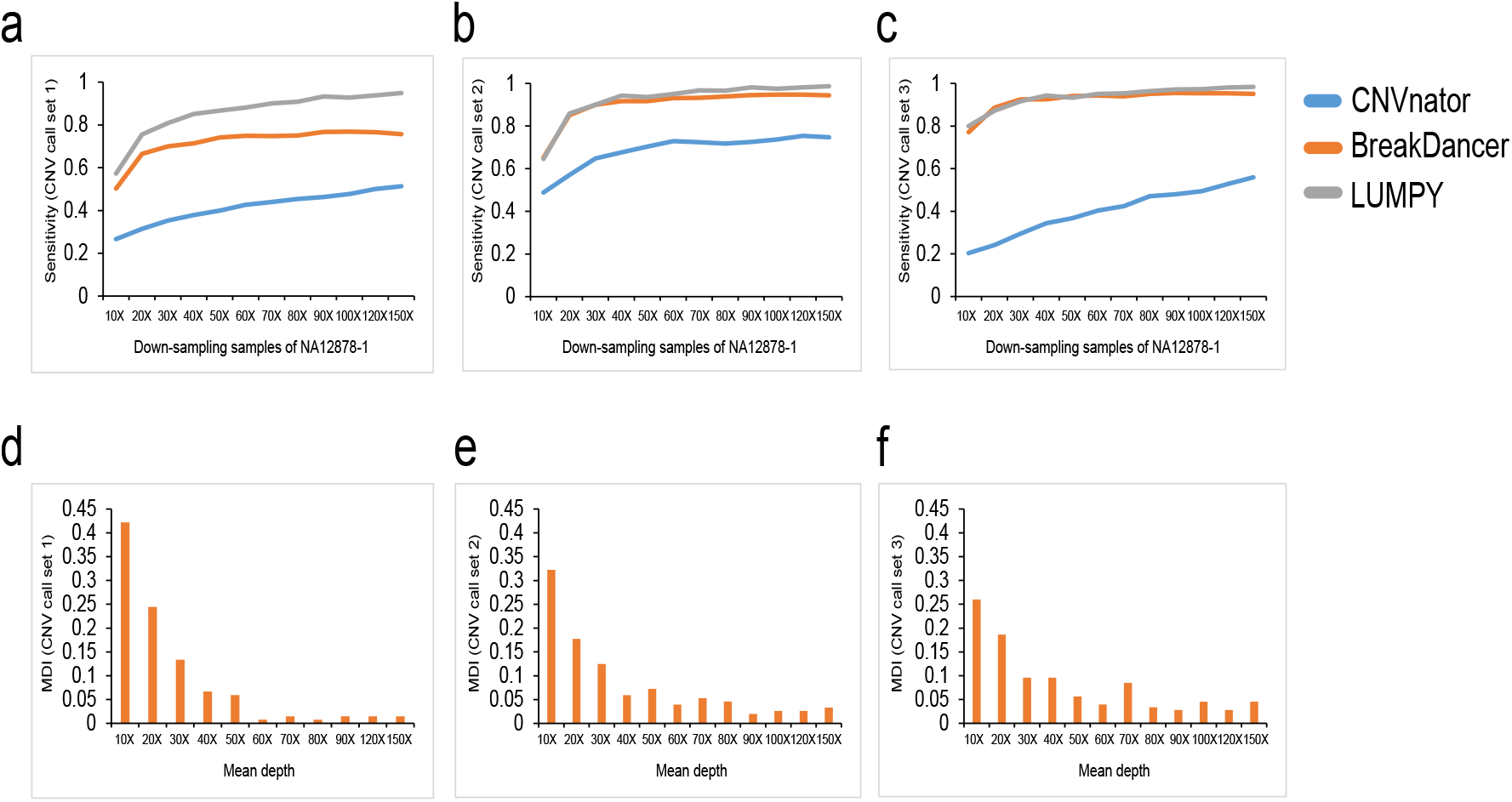
CNV detection in the 12 down-sampling samples of NA12878-1 using 3 CNV call sets

We also observed that the size of CNVs may influence the sensitivity of CNV tools. The performance of each tool varied along with the size of CNVs (Supplementary Figure 1-9). Taking the deletions in CNV call set 1 as an example, the widely-used tool CNVnator may not be suitable for CNV detection when the size of CNV is less than 1kb. When the CNV size is less than 1kb, the sensitivity significantly increased with sequencing depth from 10X to 30X but reached a plateau at a ~40X depth (Supplementary Figure 10). This indicated that when the CNV size is less than 1kb, the detection rate is greatly influenced by sequencing depth, which is less obvious when the CNV size is between 6-70kb. While, BreakDancer provided a better performance for deletion detection in all size of CNVs. These results suggested that, clinical scientists should pay more attention to the selection of CNV tools when focusing on different CNV sizes.

The selection of CNV call set and CNV detection tools may influence the sensitivity of CNV detection, making the assessment of the recommended depth for CNV detection of proband-only WGS difficult. In order to investigate the minimum requirement of mean DP for CNV detection in proband-only WGS, using 3 CNV call sets (CNV call set 1, 2, 3) and the detection results of 3 CNV tools (CNVnator, BreakDancer and LUMPY), we defined a “miss detection index” (MDI) value in this study. The MDI value for a specific mean DP is defined as the frequency when the specific mean DP shows the “lowest” sensitivity for different CNV size in a CNV call set. Without regard to selection of CNV call set and CNV detection tools, MID can be used to evaluate the recommended depth for CNV detection of proband-only WGS.

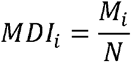

In the formula, M means the number of times when mean depth i shows the “lowest” sensitivity of CNV detection, N means the total number of times for all the depth that shows the “lowest” sensitivity of CNV detection. To obtain qualified CNV sizes in a CNV call set for evaluation, some criteria need to be fulfilled for a CNV size (Supplementary material). Detailed criteria and examples of the calculation of MDI can be found in Supplementary material.

As a result, the MDI value of a depth of 10X, 20X, 30X and 40X ranked first in the down-sampling samples of NA12878-1 (Figure 3d-f). 10X-40X account for more than 71.98% of the total depth. Taking together the sensitivity of detecting CNVs and sequencing costs, a sequencing depth of ~40X provides the best value for CNV detection, as is indicated by the trends in the sensitivity curves (Figure 3a-c).

### Depth and breadth of coverage for disease-associated genes and CNVs

Although WGS is better than WES for variation detection in patients with genetic disorders, the coverage of coding exons in key disease-associated genes of WGS has not been fully evaluated. In order to investigate the breadth of coverage of proband-only WGS for disease-associated genes, the breadth of coverage of 6 gene sets for the 12 down-sampling samples of NA12878-1 (10X-150X) and YH with increasing mean depth were evaluated. For each exons of the coding genes, we calculated the percent of exonic bases covered at more than 10X depth, which was reported to provide 95% sensitivity for heterozygous SNVs in WES [4]. None of the 12 down-sampling samples of NA12878-1 and YH covered 100% of the coding exons in the 6 gene sets except for the ACMG59 gene set (Supplementary Table 5). The results of down-sampling samples of NA12878-1 seems slightly better than down-sampling samples of YH, probably caused by the total sequencing depth of NA12878-1 (~197) and YH (~151). Across the 6 gene sets, limited range of variation was found in the down-sampling samples when the mean depth is more than 40X (Supplementary Table 5).

As for the ACMG 59 genes, we also observed a range of variation in the breadth of coverage for the 12 down-sampling samples. As such, a mean depth of more than 70X for YH and 90X for NA12878-1 covered 100% for all the ACMG 59 genes. Mean depth of 30X to 50X were most widely used for WGS [24, 25]. The proportion of genes in ACMG59 gene set covered 100% at >= 10X was 93.22%, 98.31% and 96.61% for NA12878-1_30X, NA12878-1_40X and NA12878-1_50X (with mean depths of ~30X, ~40X and ~50X). For mean depth of ~30X, ~40X and ~50X for YH, we observed that 86.44%, 93.22% and 98.31% of genes were covered 100% at >= 10X. The breadths of coverage are significantly better when the average sequencing depth is more than 40X (Figure 4a). Sites of all genes are covered more than 99.9% when the sequencing depth is ~40X. Interestingly, we also observed poorly covered RYR1 and TGFBR1 gene in this study (Figure 4a) comparing to a previously published paper for measuring sensitivity and coverage of clinical WES [4], indicating that the poor coverage of RYR1 and TGFBR1 might be caused by the features of the gene regions, not the sequencing methods used. Clinical scientists need to pay more attention to these genes when performing clinical WGS. Considering the cost of sequencing, a sequencing depth of ~40X provides the best value for the coverage of the ACMG 59 gene set, as is indicated by the trends in breadth of coverage value of the ACMG 59 genes. We observed similar patterns for down-sampling samples of YH (Supplementary Table 6). We also examined the percentage of genes at >= 20X coverage in the ACMG 59 gene set, which could provide 99% sensitivity for heterozygous SNVs [4]. 81.36% and 59.32% of genes were covered 100% for NA12878-1 and YH when the mean DP is ~40X.

**Figure 4.**
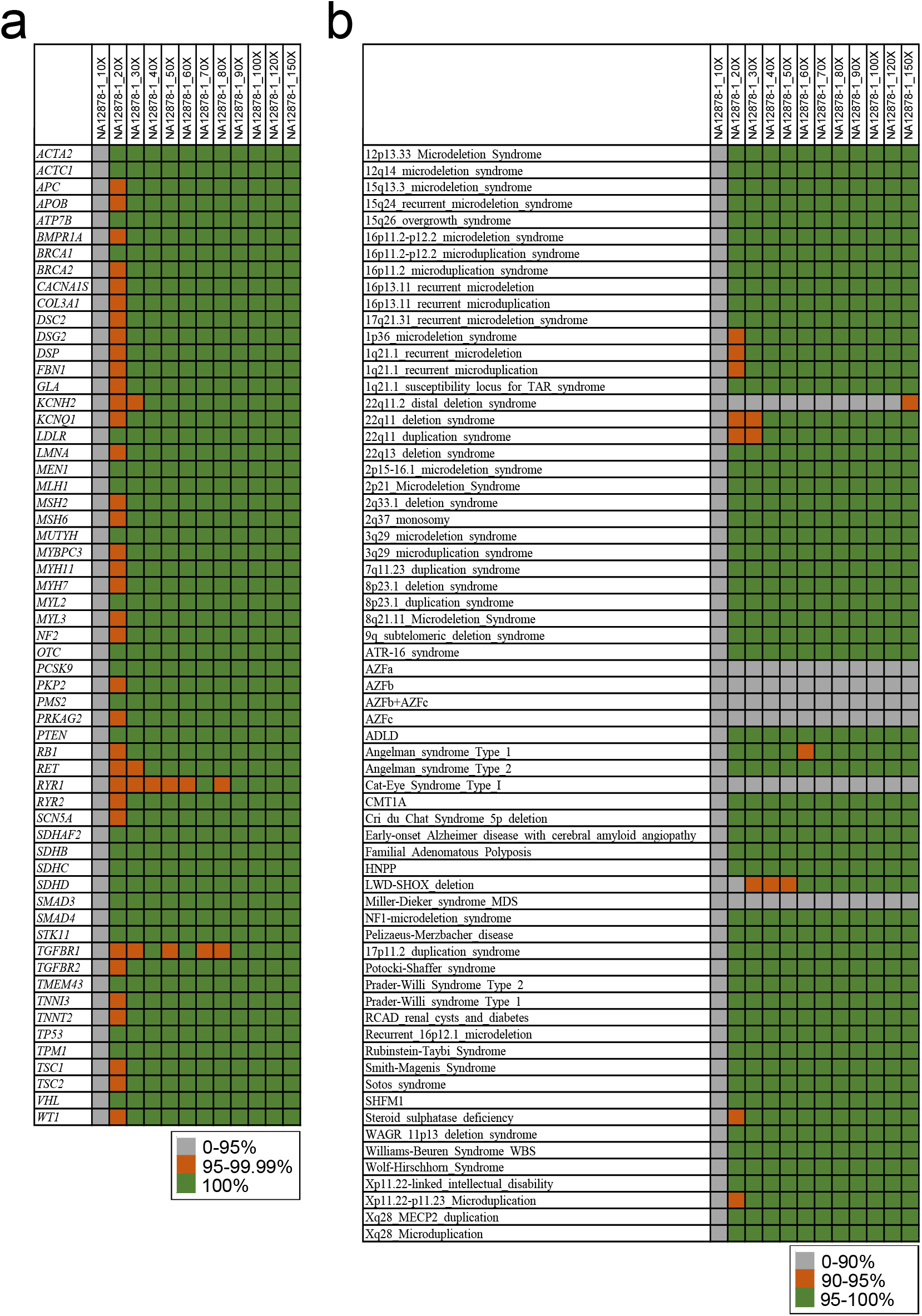
Depth and breadth of coverage for disease-associated genes and CNVs

CNVs are another major part of human genetic variation, which are often found to be associated with human diseases. For the CNVs in the DECIPHER database, most CNVs can be well covered (more than 95% coverage) at >10X depth when the sequencing depth is 40X-50X for NA12878-1 (Figure 4b). Clinical scientists should pay more attention to 22q11.2 distal deletion syndrome, Cat-Eye Syndrome Type I and Miller-Dieker syndrome (MDS), which showed a coverage of 88.96% and 4.54% even with sequencing depth of 100X. Similar patterns were also observed for down-sampling samples of YH (Supplementary Table 2).

### Sensitivity and PPV among trios

The purpose of the analysis of trios, is to test the sensitivity and PPV when taking advantage of the family-based trio information in clinical WGS. Although the sequencing depth of the Chinese trio is more than 100, here we used down-sampling samples with a mean depth of 44.00X, 43.77X and 43.05X for NA24631, NA24694 and NA24695. The reason why we concentrate on the depth of ~40X is that, ~40X depth is the most widely used and recommended depth for WGS, which is also consistent with some of our previous results, especially for CNV detection.

In this study, we took advantage of the family-based trio design (Figure 5a) to calculate the sensitivity and PPV in high confidence regions (NISTv3.3.2/GRCh37) [38]. The analysis was restricted to variants with DP >= 10X and GQ >= 20 (Figure 5a). We focused on the loci where one parent was homozygous for the alt allele and the other was homozygous for the reference allele. For these variants, the offspring should be heterozygous. This could provide a new “gold standard set” for NA24631 (the offspring). Comparing to the high-confidence calls of NA24631 provided by NIST (the high-confidence call set), the new gold standard set could be used to test the sensitivity and PPV for trio-based WGS. Figure 5b showed the results of the sensitivity and PPV of the “gold standard set” and the high-confidence call set for NA24631. As a result, the sensitivity of the “gold standard set” for SNP and indel detection was >99.48% and >96.36% respectively, and PPVs were 99.86% and 97.93%. Trio-based analysis showed great improvement for the PPV of indel detection (Figure 5b), PPV for indel detection improved from 89.68% (using the high-confidence call set) to 97.93% (using the “gold standard set”).

**Figure 5.**
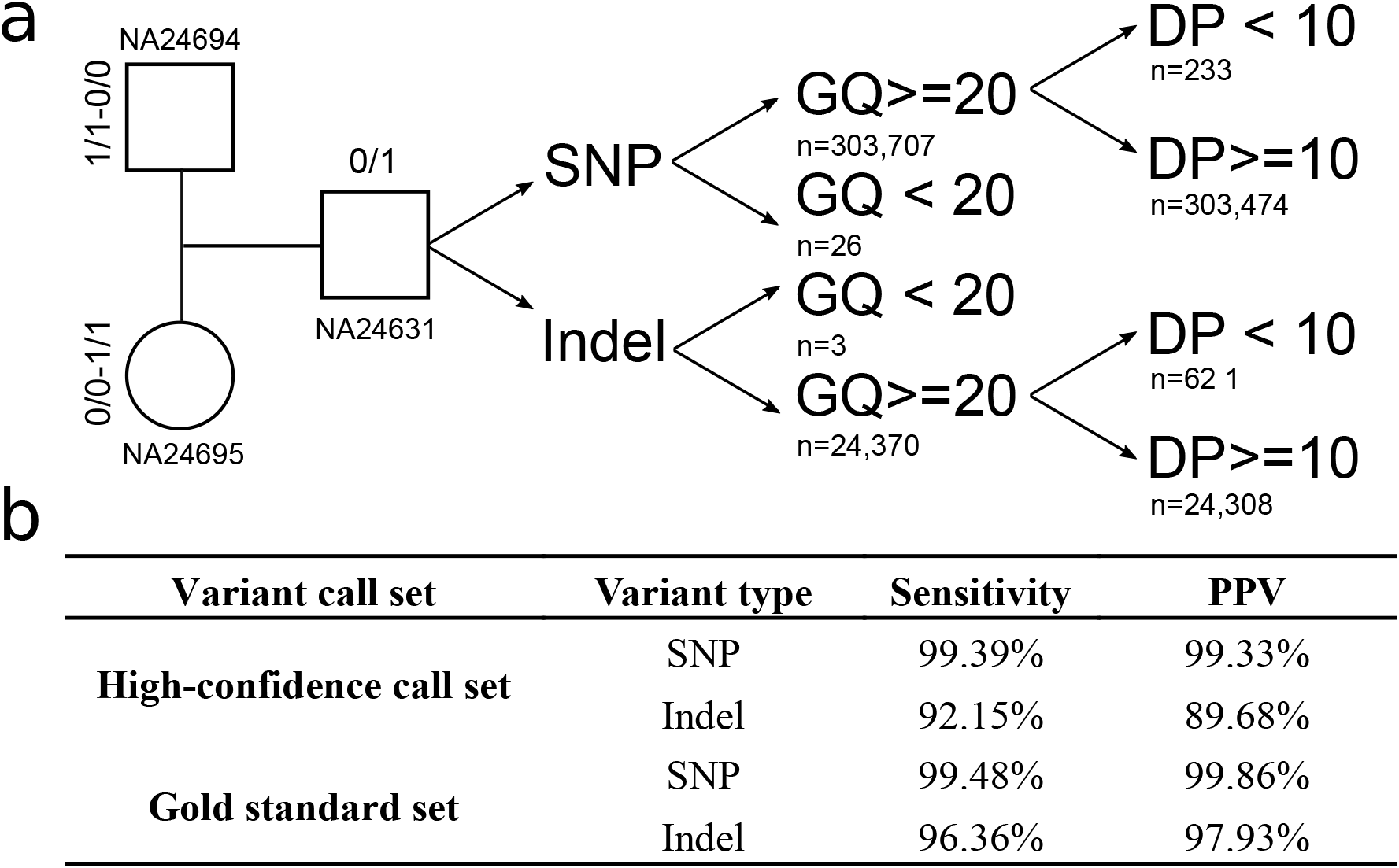
Variants analysis in the Chinese trios

### WGS and WES

Here, we evaluated the performance of WES (MGIEasy Exome FS Library Prep Set) and WGS (MGIEasy PCR-Free DNA Library Prep Set) on DNA samples of NA12878. NA12878-1_40X and NA12878-2_120X were used for evaluation. NA12878-1_40X was the down-sampling sample of NA12878-1 with a mean DP of 40X. NA12878-2_120X was an down-sampling sample of NA12878-2 with a mean DP of 120X, which is typical and current standard for clinical WES [49]. Two quality parameters for variation detection (DP and GQ), sensitivity for SNV/indels detection, breadth of coverage of the list of 8,394 putative disease-associated genes and CNVs were compared in this part.

DP and GQ are two main parameters assessing quality of variant calls, which are often used to filter out variants with erroneous variant calls [30, 33]. First, we investigated the GQ and DP distribution of NA12878-1_40X and NA12878-2_120X in the regions of the human genome covered by WES (59,082,036 bp). We found similar results to those obtained in a previous WES study [8]. The distribution of DP for the variants ranged more widely in NA12878-2_120X than in NA12878-1_40X (Supplementary Figure 11), with a median at 94X depth but a mode at 63X and 58X for SNPs and indels, indicating low levels of coverage for a substantial proportion of variants. While the distribution of DP was normal-like for NA12878-1_40X, with a median at 42X and coinciding mode at 41X for both SNPs and indels (Supplementary Figure 11). The vast majority of variants called by NA12878-2_120X had a GQ close to 100, and fluctuated along with the GQ scores. The distribution of GQ for variants in NA12878-1_40X showed a mode in low GQ area, probably caused by insufficient variant calls in the WES regions.

For the detection of SNVs and indels, “true positive” calls were further restricted to the regions of the human genome covered by WES (59,082,036 bp). A total of 44,726 variants were used for evaluation. DP and GQ filtering were not used in this part. In general, the sensitivity and PPV of WGS (NA12878-1_40X) is higher than WES (NA12878-2_120X) for the 44,726 variants (Supplementary Table 7), except for the PPV for homozygous indel detection. For 90.39% of gold SNPs, WES and WGS yielded the same genotype. More than 63.41% of these concordant SNVs were identified as heterozygous, which was similar to those obtained in previous WES studies [4, 15, 25].

Then, we investigated the breadth of coverage of NA12878-1_40X and NA12878-2_120X in the 8,394 putative disease-associated genes and CNVs (DECIPHER database). As a result, WGS showed better coverage of both putative disease-associated genes and CNVs. More than 99.77% sites of the exon regions of the 8,394 putative disease-associated genes were covered with a depth of >=10 for NA12878-1_40X, while NA12878-2_120X covered 99.46% of the exon regions. NA12878-2_120X was poorly covered for CNVs in the DECIPHER database (Supplementary Table 8). More than 69.69% of the CNVs showed a coverage of less than 10%.

### Analysis of 8 clinical cases with known disease-causing variants

In this study, samples of 8 clinical cases with known variants of various types were recruited and reanalyzed using our WGS pipeline. All the 12 variants were validated previously by methods other than MPS technology. Focusing on SNVs, indels and CNVs, we applied our method to 8 clinical cases using singleton WGS. Variants were manually assessed for quality and interpreted according to the American College of Medical Genetics and Genomics (ACMG) guidelines for variant classification [27, 28]. All the previously validated variants (Supplementary Table 9) were successfully detected by our WGS pipeline, which further demonstrated the sensitivity of the method.

Based the sequencing DP, the average cost for the WGS approach was ~$490 per sample for the 8 clinical cases in current study. The overall cost (from DNA extraction to reporting) for a single case with a mean sequencing depth of ~40X is approximately $600, including ~$280 for chemicals (DNA extraction, library construction and sequencing), ~$220 for labor, and ~$100 for depreciation expenses.

## Discussion

The emergence of MPS technology makes multigene sequencing, exome level sequencing, and even genome level sequencing possible, which has been more and more widely used in clinical diagnosis for genetic disease. In this study, we performed a comprehensive analysis of sensitivity and coverage of clinical WGS as a diagnostic test for genetic disorders. First, we analyzed the sensitivity and PPV value of high-confidence SNPs/indels, and sensitivity of CNV detection using down-sampled NA12878 and YH. A new MDI value was defined for the evaluation of CNV detection; then we investigated the depth and breadth of coverage for disease-associated genes and CNVs in down-sampling samples of NA12878 and YH. A new gene set and a CNV call set were generated during this process; then we compared the performance of WES and WGS on DNA samples of NA12878; we also tested the sensitivity and PPV of variant calls when taking advantage of the family-based trio design in clinical WGS; finally, we analyzed 8 clinical cases with known disease causing variants using our WGS pipeline.

The results suggested that, WGS can be used in the detection of SNVs/indels/CNVs with high sensitivity, the current standard of a mean depth of ~40X may be a cost-effective sequencing depth for SNV/indel detection and the identification of most CNVs, WGS is bound to be widely used and become a routine part of clinical care in the near future.

During the analysis of the PPV of indels in the high confidence regions, we detected a slightly decline for heterozygous indels in the 12 down-sampling samples of NA12878 with the increase of DP. One reasonable explanation is that, with the increase of depth, both of the true positives and false positives increases. It turned out that the false positives increased a little bit faster than the true positives, as a result, the PPV declined. Massive scale p population based polymorphism database and further filtering of the variants may be useful to solve this problem.

In addition to SNVs and indels, another major part of human genetic variation is copy number variation. According to the clinical requirements, choosing the suitable methods and tools for accurate and reliable detection of CNVs is important for clinical diagnostics. WGS can detect nearly all known genetic variations. However, our results indicated that, although CNV calling is reliable with increasing DP, the performance of CNV tools varied a lot. Finding the right tool for CNV detection is difficult for clinical scientists. Our results suggested that read pair methods (BreakDancer in particular) were the best performing approaches for the identification of deletions of more than 1 kb. What is more, although some “gold standard” CNV call set is widely used in published papers [45–47], lacking validation of various methods, some CNVs may be false positives with wrong or low resolution boundaries. Factors (such as CNV size and the selected “gold standard” set) may also influence the sensitivity of CNV detection, making it difficult to determine the sufficient DP for CNV detection. In this study, we introduced the concept of MDI to solve this problem. Finally, we found that the current standard of a mean depth of 40X may be sufficient for the identification of most CNVs. However, there are limitations of our analysis: we only sampled a small number of CNV callers (CNVnator, BreakDancer and LUMPY). In real clinical setting, the application of more than one CNV calling algorithm should be considered to improve the sensitivity of CNV detection. Analyzing sensitivity of CNV detection with all available CNV tools would be an interesting research topic. In addition, we found that additional coverage is associated with the overall increases of sensitivity for CNV detection, however, this is less obvious as the CNV size is more than 100kb, which has also been found in another published paper [50].

There are no perfect “gold standard” CNV dataset for benchmarking. So, in this study we compiled a list of 2,022 “likely true positive” from the 3 most commonly used CNV call sets of NA12878 from published papers [45–47] for benchmarking (Supplementary Table 10), including 1,912 deletions and 110 duplications. This new set is a combination of the 3 CNV call sets after evaluation. For this new CNV call set, we defined true positives as the variants were detected by at least one CNV tool (CNVnator, BreakDancer and LUMPY) with more than 50% reciprocal overlap and confirmed by visualization of copy ratio though an in-house script. This new set is an ideal “gold standard” CNV call set of NA12878 for clinical WGS benchmarking.

Turn-around time and cost of WGS are two key point for clinical application of WGS. The entire workflow of this method lasts approximately 11-12 days from the recruitment of sample to clinical reporting for one sample. BGI produced MegaBOLT (MegaBOLT bioinformatics analysis accelerator) along with the sequence platform MGISEQ-2000, which is an MGI self-developed and MPS-concentrated hardware accelerating system for bioinformatics analysis. MegaBOLT supports the analysis of WGS and WES, and is 20 times faster than the traditional GATK approach, which can be used to shorten the bioinformatics process. Along with the development of automated diagnosis tools [51, 52], which could be used to prioritize patient phenotypes and expedite genetic disease diagnosis, the turn-around time of WGS could be further reduced. The overall cost, including chemicals, labor, and depreciation expenses for the WGS approach was $600 per sample (~40X depth). Sequencing accounted for nearly half of the total cost. In general, variant calling is more reliable with increasing DP. However, there are detection ceiling for some genes and/or regions (such as regions related to Miller-Dieker syndrome), which cannot be solved by increasing the sequencing depth. The cost and sensitivity of WGS need to be balanced. Our results suggested that the current standard of a mean depth of 40X may be sufficient for the identification of most SNVs and CNVs. The reduce of the cost and turn-around time would further improve the clinical application of WGS.

## Conclusion

In summary, the successful application of WGS as a diagnostic test for genetic disorders in real clinical setting requires comprehensive assessment of the depth and breadth of coverage, and sensitivity of WGS. In this study, we observed variation in the detection of SNV/indel/CNV, and substantial variation in coverage of medically implicated genes and CNVs. In real clinical setting, clinical scientists should know the range of sensitivity and PPV for different classes of variants for a particular WGS pipeline, which would be useful when interpreting and delivering clinical reports. We believe that WGS is likely to change the way of clinical diagnosis of rare and undiagnosed diseases in the near future.

## Declarations

### Ethics approval and consent to participate

This study and all the protocols were approved by the ethics committee of BGI (NO. BGI-IRB19143).

### Consent for publication

Not applicable.

## Availability of data and material

The data that support the findings of this study have been deposited in the CNSA (https://db.cngb.org/cnsa/) of CNGBdb with accession code CNP0000813. The data of the 8 clinical cases generated and analyzed during the current study is not publicly available as they are patient samples and sharing them could compromise research participant privacy. The datasets of the GIAB samples used and analyzed during the current study are available from the corresponding author on reasonable request.

## Authors’ contributions

ZYP, JS, and YS designed the research. YS wrote the first draft of the article. CNF, LJS, XDW, ZYY, ZPX, CNS, HYZ, WD, YL, YQL and JS designed and performed the experiments. FXL, YSW, ZHF, RH, ZHW, JGP and ZYP performed data analysis. HH provided patient specimens and conducted histopathological examinations. ZYP, JS, YS, FXL, CNF, YSW and LJS contributed to drafting and revising the manuscript. All authors reviewed the manuscript.

## Acknowledgments

We thank all the blood donors for their invaluable contribution to this study.

---

**Table 1** Sample information

**Additional files**

**Additional file 1:** Supplementary Tables

**Additional file 1:** Supplementary material

